# Improving oral dissolution kinetics of weakly basic vodobatinib via slurry conversion to an amorphous drug-polymer salt

**DOI:** 10.64898/2026.06.26.734800

**Authors:** Luke D. DeLion, Sophia R. Dasaro, Mojhdeh Baghbanbashi, Dmitry Y. Zemlyanov, Kurt Ristroph

## Abstract

Vodobatinib (VBN) is a weakly basic (pK_a_ ≈ 2.3), anticancer treatment with poor enteric solubility and low oral bioavailability. This study demonstrates how an emerging polymeric amorphization technique, slurry conversion, can yield amorphous drug-polymer salts with enhanced dissolution rates. The technique had not previously been applied to a weakly basic drug, so design rules for this class of active were unknown. Two acidic polymers, poly(styrene sulfonic acid) (PSSA) and poly(acrylic acid) (PAA), were individually evaluated for salt formation with VBN. Formulation involved blending the drug and polymer in a 1:2 (v/v) ratio of a protic liquid to solvent and a 1:9 (w/w) ratio of solid to solvent. Design rules for effective combinations of solvents and protic liquids were developed and optimized to thread the needle between dissolution of all species and acid-base interactions, both of which were required to form amorphous salts. Drug loadings of 10%, 20%, and 40% by mass were tested. X-ray photoelectron spectroscopy was employed to evaluate protonation of the quinoline nitrogen atoms on VBN, a key indicator of successful salt formation. Powder X-ray diffraction was used to confirm that the resulting slurry contained amorphous VBN, and ^1^H NMR spectroscopy indicated residual solvent remained after drying, which remains an area for improvement. In dissolution kinetics tests in FeSSIF, the lead drug-polymer salt formulation achieved a concentration of dissolved VBN up to 140 µg/mL, an improvement of >35-fold compared to <4 µg/mL (LLD) for crystalline VBN. These results demonstrate that slurry conversion is a viable polymeric amorphization technique even for weakly basic drugs.

**Graphical abstract:** 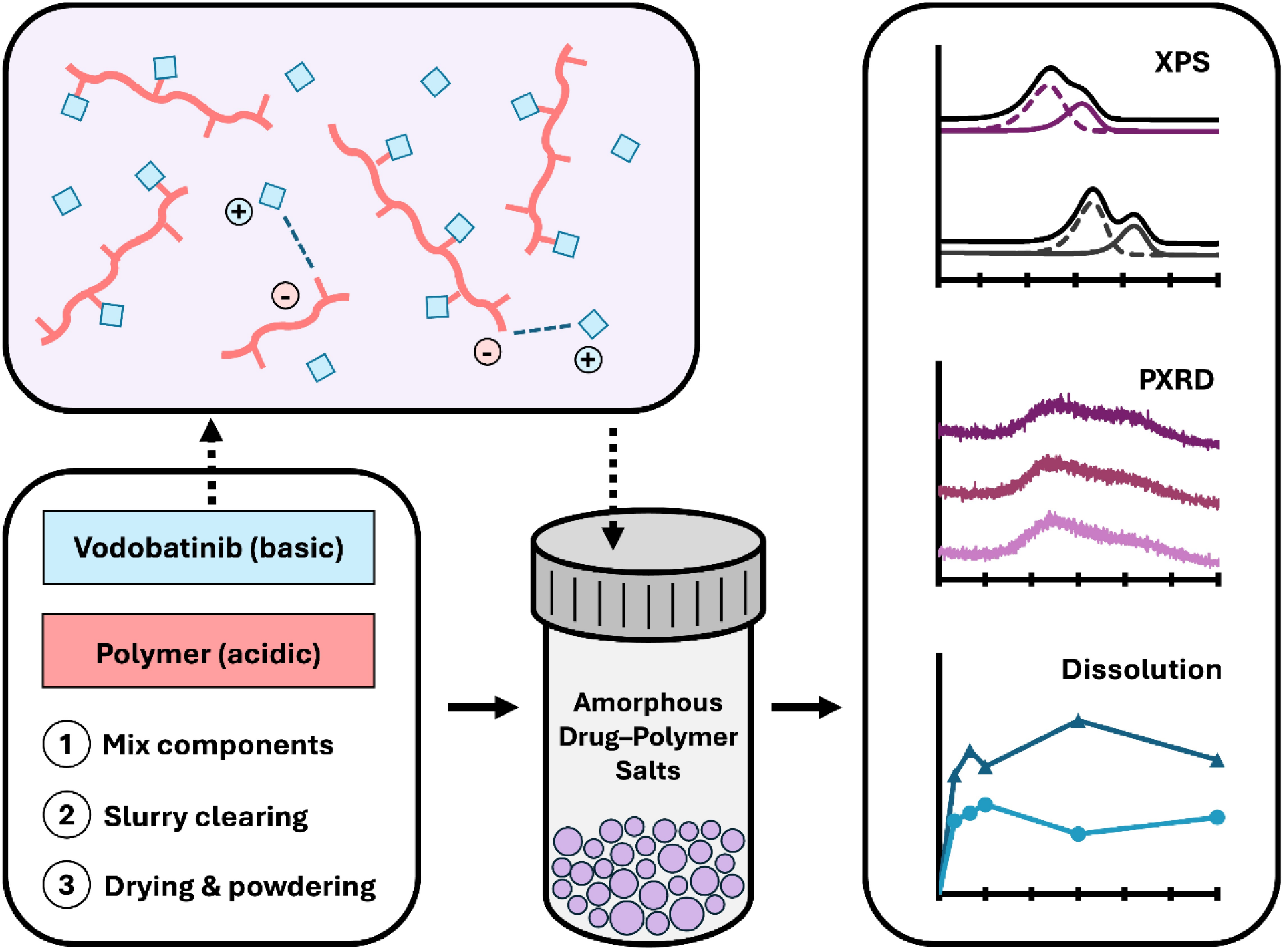

## 1. Introduction

Small-molecule drugs account for the majority of FDA-approved therapeutics and continue to dominate current medical treatments, but their clinical success is often limited by low oral bioavailability driven by poor solubility and high hydrophobicity [1-4]. After oral administration, enteric dissolution of the active pharmaceutical ingredient (API) is essential for maximizing circulatory distribution and minimizing dosing burden [5, 6]. With an estimated 90% of developmental drug candidates exhibiting poor aqueous solubility, amorphization has become a common strategy for improving API druggability [6-10]. Amorphous compounds have no long-range molecular structure, resulting in a high energy state which facilitates increased water solubility and faster dissolution rates [11, 12]. Despite these advantages, amorphous products are prone to recrystallization and hygroscopic instability, thus selecting a proper manufacturing method and formulation that preserves the API in an amorphous state for long periods of time is crucial [13, 14].

Typical formulation methods for manufacturing amorphous APIs consist of hot melt extrusion and spray drying [11, 15, 16]. While these approaches are well-suited for large-scale manufacturing, limitations such as high equipment costs, elevated processing temperatures, and excipient constraints have motivated the exploration of alternative methodologies. One emerging approach, referred to as slurry conversion, has shown promise as a low-cost and comparably lower-energy strategy to enhance oral bioavailability of APIs [17-20]. This technique is founded on the principles of salt formation, specifically the acid–base interactions between ionizable functional groups on the API and an oppositely-charged counterion [16]. For effective salt formation, the ΔpKa between the interacting species should generally exceed 2–3 units [21]. In practice, slurry conversion involves dispersing a drug and selected counterion in a solvent, followed by solvent evaporation at substantially lower temperatures than those required for spray drying [17]. The resulting material is an amorphous solid with improved biopharmaceutical performance [18, 19]. To date, this approach has been applied to both solid and liquid dosage forms using small inorganic or organic counterions [19, 22].

Recent efforts in slurry conversion have increasingly focused on the use of water-soluble polyelectrolytes as counterions instead of small molecules. Polyelectrolytes exhibit high hydrophilicity which can enhance aqueous wettability and apparent solubility [19]. For example, Neusaenger *et al*. found that forming salts by protonating basic drugs with poly(acrylic acid) (PAA) increased dissolution rates up to 6 times that of other formulation methods [17]. In addition to improved dissolution, drug–polymer ionic interactions are stronger than the intermolecular forces present in neutral systems, often resulting in elevated glass transition temperatures and improved stability under thermal and humid stress compared to melt-extruded formulations [18]. Another key advantage of slurry conversion lies in the wide range of acidic polymers capable of forming salts with basic drugs [17, 19]. However, this versatility is not universal; weakly basic APIs often fail to form salts with the polyanions commonly employed in this approach, limiting its applicability to this important class of therapeutics [15].

In this study, we extend the slurry conversion strategy to a particularly challenging weakly basic API, vodobatinib (VBN), with the goal of identifying conditions that enable successful salt formation. VBN is a poorly soluble promising treatment for chronic myeloid leukemia, with clinical trials for Lewy body dementia and Parkinson’s disease currently under review [23, 24]. Since VBN is a weak base with an estimated pK_a_ of 2.3, this API is a complex candidate for salt formation [25]. Consequently, there exists a limited set of polymeric counterions possess sufficient acidity and hydrophilicity to facilitate effective complexation with VBN. Poly(styrene sulfonic acid) (PSSA), with a pKa of −1.5 [12] and high aqueous solubility, satisfies these criteria and is already widely employed in biomedical applications [26, 27]. Accordingly, PSSA was selected to promote VBN protonation, while PAA (pKa = 4.5), a substantially weaker acid with historical uses in drug-polymer slurry formation, was evaluated as a negative control.

The results presented herein demonstrate that slurry conversion successfully produces VBN–PSSA salts, across a range of drug mass loadings, with amorphous character and enhanced dissolution rates. Each drug–polymer salt was confirmed to be amorphous using powder X-ray diffraction, and salt formation was verified with X-ray photoelectron spectroscopy. After slurry evaporation, residual solvents were qualitatively evaluated using ^1^H NMR spectroscopy. The results of this analysis identified that residual solvents were present in all salts, necessitating a more robust approach to final solvent removal. All successful salt formulations exhibited faster dissolution kinetics compared to both the non-salt formulations and crystalline VBN in fed state simulated intestinal fluid. Collectively, these results demonstrate the feasibility of improving oral bioavailability for weakly basic APIs, which expands the application space for the slurry conversion formulation technique.

## 2. Materials and methods

### 2.1 Materials

Poly(acrylic acid) (PAA, M_v_ ∼450,000) and poly(styrene sulfonic acid) (PSSA, M_w_ ∼75,000, dissolved in water at 30% w/v) were purchased from Sigma-Aldrich (St. Louis, MO, USA). Vodobatinib (VBN) was provided by Sun Pharma Advanced Co (Mumbai, India). Ethanol, acetonitrile (ACN), and tetrahydrofuran (THF) were purchased from Thermo Fisher Scientific (Waltham, MA, USA). Fed state simulated intestinal fluid (FeSSIF) and fasted state simulated gastric fluid (FaSSGF) powder and buffer were purchased from Biorelevant.com (London, UK). All materials were used as received.

### 2.2 Drug-polymer salt preparation

Amorphous salts were prepared as an adapted procedure from Yao *et al.* [19]. Crystalline VBN was weighed and sonicated with an organic solvent (THF or ACN). An aqueous solution of PSSA was then added to form a slurry with a 1:9 (w/w) solid/solvent ratio and a 1:2 water/organic solvent ratio. Formulations were prepared at 10, 20, and 40% (w/w) drug loading, such that the total drug–polymer mass was 200 mg. Each mixture was magnetically stirred for up to 2 hours in a silicone oil bath at 60 or 75 °C, depending on solvent boiling points, and dried under a heated vacuum for 1 day. The resulting salt was ground into particles with a ceramic mortar and pestle under liquid nitrogen, dried in a vacuum desiccator overnight, and further powdered using an agate mortar and pestle. To evaluate the potential advantages of salt formation, a theorized non-salt formulation with PAA was prepared. VBN was dissolved in THF, then PAA dissolved in ethanol was added. The same mixing and drying methods were followed. Due to the hygroscopicity of both polymers, all samples were stored in a vacuum desiccator to prevent excess moisture uptake and particle adhesion. An overview of the relevant materials and workflow employed here is provided as **Figure 1**.

**Figure 1:**
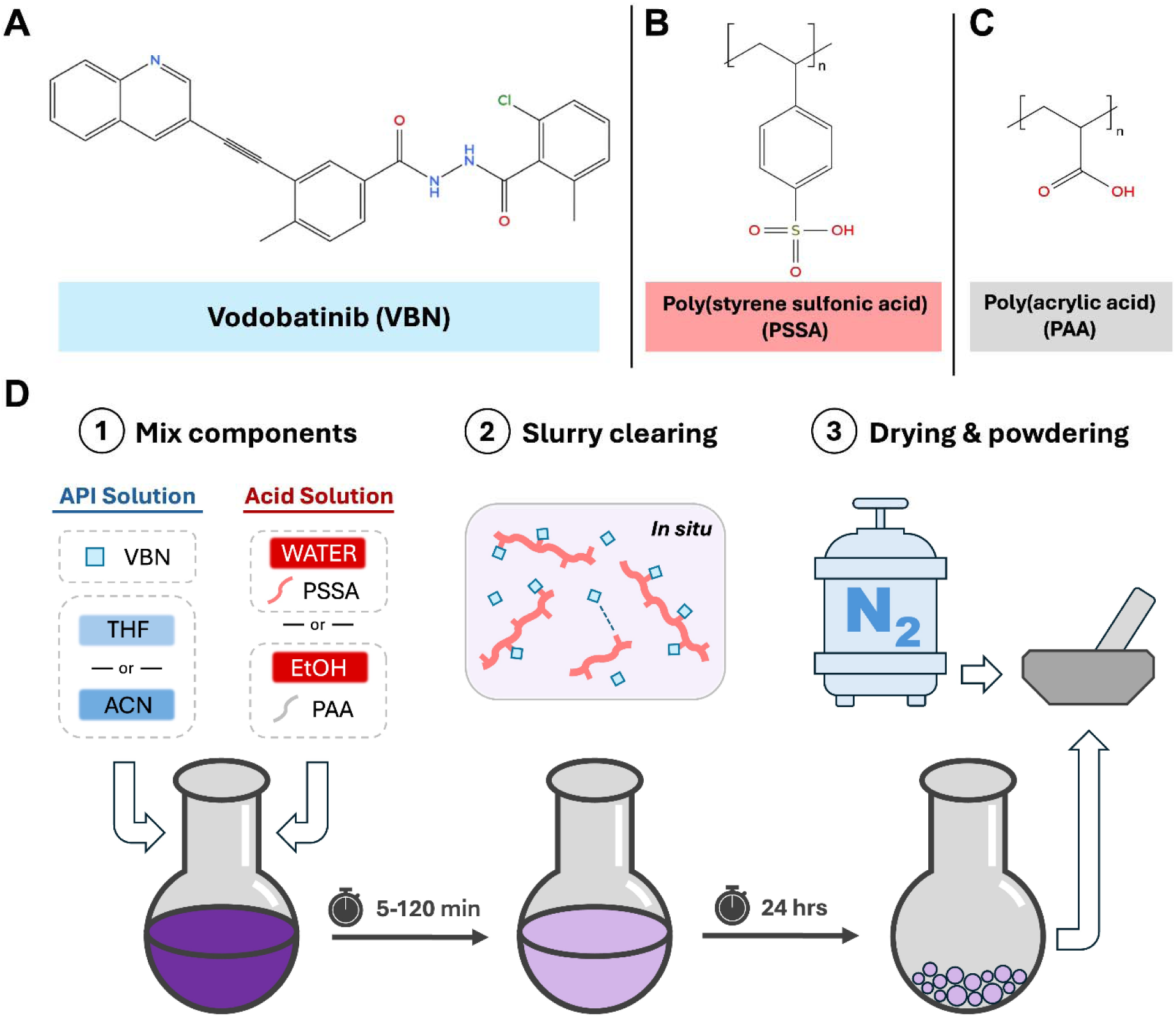
(A-C) Chemical structures for key slurry components including (A) vodobatinib (VBN), (B) poly(styrene sulfonic acid), (PSSA), and (C) poly(acrylic acid), (PAA). The experimental workflow is presented in panel (D).

### 2.3 Powder X-ray diffraction (pXRD)

Powder X-ray diffraction was performed in focusing mode on a Panalytical Empyrean X-ray diffractometer equipped with Bragg-Brentano HD optics, a PixCel3D Medipix detector, and a Cu Kα X-ray source (λ = 1.54178). Powdered samples were measured at ambient temperature with a tube power of 45 kV and 40 mA. A rotating metal stage with a circular sample area of 16 mm diameter and 2 mm deep was used for analysis. Anti-scatter slits, divergence slits, and the mask were selected based on the scanning range and stage area. Each powder was scanned 6 times between 5 and 55° (2θ) with a 0.013° step size using Panalytical Data Collector software [28, 29]. The sum of 6 scans was collected in HighScore and exported to Excel for visualization.

### 2.4 X-ray photoelectron spectroscopy (XPS)

XPS measurements were performed using a Kratos Axis Ultra DLD spectrometer with monochromatic Al Kα radiation (1486.6 eV). Survey spectra were acquired at a pass energy of 160 eV with a step size of 1 eV. High-resolution N 1s and C 1s spectra were collected at a pass energy of 20 eV with a step size of 0.05 eV. A commercial Kratos charge neutralizer was used to improve spectral quality.

Data were analyzed using CasaXPS software. For charge correction, the C 1s component corresponding to C–C bonds was set to a binding energy of 284.8 eV. Peak fitting of the C 1s and N 1s spectra was performed using Gaussian–Lorentzian line shapes with a 30% Lorentzian contribution (GL(30)) and a Shirley or linear background subtraction.

### 2.5 Dissolution kinetics testing in simulated gastrointestinal fluids

*In vitro* dissolution kinetics were evaluated in fed-state simulated intestinal fluid (FeSSIF) at 37 °C. Media was prepared by following given product instructions. Aliquots containing each VBN-PSSA or VBN-PAA product were prepared in triplicate, such that a test concentration of 1 mg/mL VBN was achieved after addition of the release media, as reported previously [30]. Samples at 10% and 20% drug loading were suspended in 100 µL of MQ for 15 minutes to mimic powder dispersion in a sachet. The gastric environment was simulated by adding 10 µL of 11x FaSSGF to the solution and incubating for 15 minutes at 37 °C. 1.12x FeSSIF was then added such that the total volume was 1 mL and with the following final conditions: 0.1x FaSSGF, 1x FeSSIF, 1 mg/mL VBN. Samples with 40% drug loading were prepared at double the volume to enable more accurate mass measurements. Drug dissolution was measured at 20 min, 40 min, 1 hr, 3 hr, and 6 hr. At each timepoint, samples were centrifuged at 30,000 rpm for 15 minutes and the supernatants were recovered (900 µL for 10, 20% samples and 1.8 mL for 40% samples). Dissolution was measured using UV-Vis spectrometry at a wavelength of 317 nm (selected based on a maximum signal from the drug and minimum contribution from the polymer) with necessary dilutions, and volume calculations were performed to accurately compare each sample using a standard curve of known VBN concentrations. The UV–Vis spectrometer was blanked with the release medium to minimize background absorbance. To correct for polymer interference at 317 nm, the polymer content of each slurry sample was estimated from the nominal formulation composition. For example, a 10 mg slurry sample containing 10 wt% drug was assumed to contain 9 mg PSSA. The corresponding PSSA absorbance was calculated from the PSSA calibration curve and subtracted from the measured sample absorbance to determine the drug contribution.

### 2.6 Nuclear Magnetic Resonance (NMR)

Proton (^1^H) NMR spectra were acquired at ambient temperature using a Bruker AVANCE III HD 500 MHz (AV500HD) spectrometer equipped with a 5 mm BBFO Z-gradient Prodigy cryoprobe and dual RF channels with waveform generation capability. Data processing and spectral analysis were carried out with Bruker TopSpin software (v4.3.0).

## 3. Results & Discussion

### 3.1 Production of drug-polymer slurries

The formation of a successful drug-polymer slurry can be preliminarily determined by observation of the material throughout processing, particularly looking at whether the solution turns clear after combining the materials. The initial formulation screen was designed to examine three critical formulation parameters – polymer type, drug loading, and solvent selection – as prior work by Yao *et al*. has established that selection of an appropriate solvent, among the other parameters, is necessary to ensure the final product retains amorphous character [19]. To ensure that the drug-polyanion species are maximally charged and can interact favorably, an appropriate solvent environment must be selected. The mixed solvent system must meet three criteria: (1) one solvent must be a good solvent for the drug; (2) the other must be a good solvent for the polymer; and (3) at least one of the solvents must be sufficiently protic such that the mixed solvent can facilitate charge transfer between the two ionizable species.

Isolation of the three examined solvent quality conditions was established by first considering the balance required to enable the molecular dissolution of both components that is required for salt formation. Specifically, VBN will dissolve in both THF and ACN at room temperature at low enough concentrations but exhibits poor water solubility that leads to rapid precipitation in the mixed water-solvent environments generated upon addition of PSSA, which is provided in water. PSSA will precipitate slightly in THF, but not ACN. In experiments where PSSA was utilized as the polymer, both THF and ACN were screened. PAA and VBN are both readily solubilized in the mixed THF-ethanol system and thus other formulations were not considered.

To determine whether complete dissolution of the polyanion and drug occurred, a “clearing” phenomenon was used as a visual indicator. This metric was adapted from previous studies by Neusaenger *et al*. investigating a lumefantrine-PAA system. For the lumefantrine-PAA system, conversion yielded an abrupt transition from a liquid slurry-like phase into a clear suspension [15, 17]. That prior work also found that salts containing very weakly basic, poorly-soluble APIs such as albendazole and mebendazole exhibited non-amorphous behavior when clarity was not reached [15, 17]. Here, we observe a similar transition for seven of the nine tested formulations. Within 5 minutes of achieving optical clarity, though, two of the VBN-PSSA acetonitrile/water formulations became turbid upon cooling. This behavior has not been previously reported and may be attributable to a reduction in transient solubility as temperature drops. These two ‘transient’ drug-polyanion slurry formulation candidates were subjected to the same physicochemical characterization as the other five lead formulations. Comparative analysis was conducted to determine whether sustained solution clarity during cooling is required to enable successful slurry formation, or if transient clarity at elevated temperature is sufficient to predict process viability. This phenomenon is important to understand, as the control formulations containing PAA, a polyanion with a pKa that should not enable salt formation, also reached clarity. The results of the initial formulation screen are presented in **Figure 2**.

**Figure 2:**
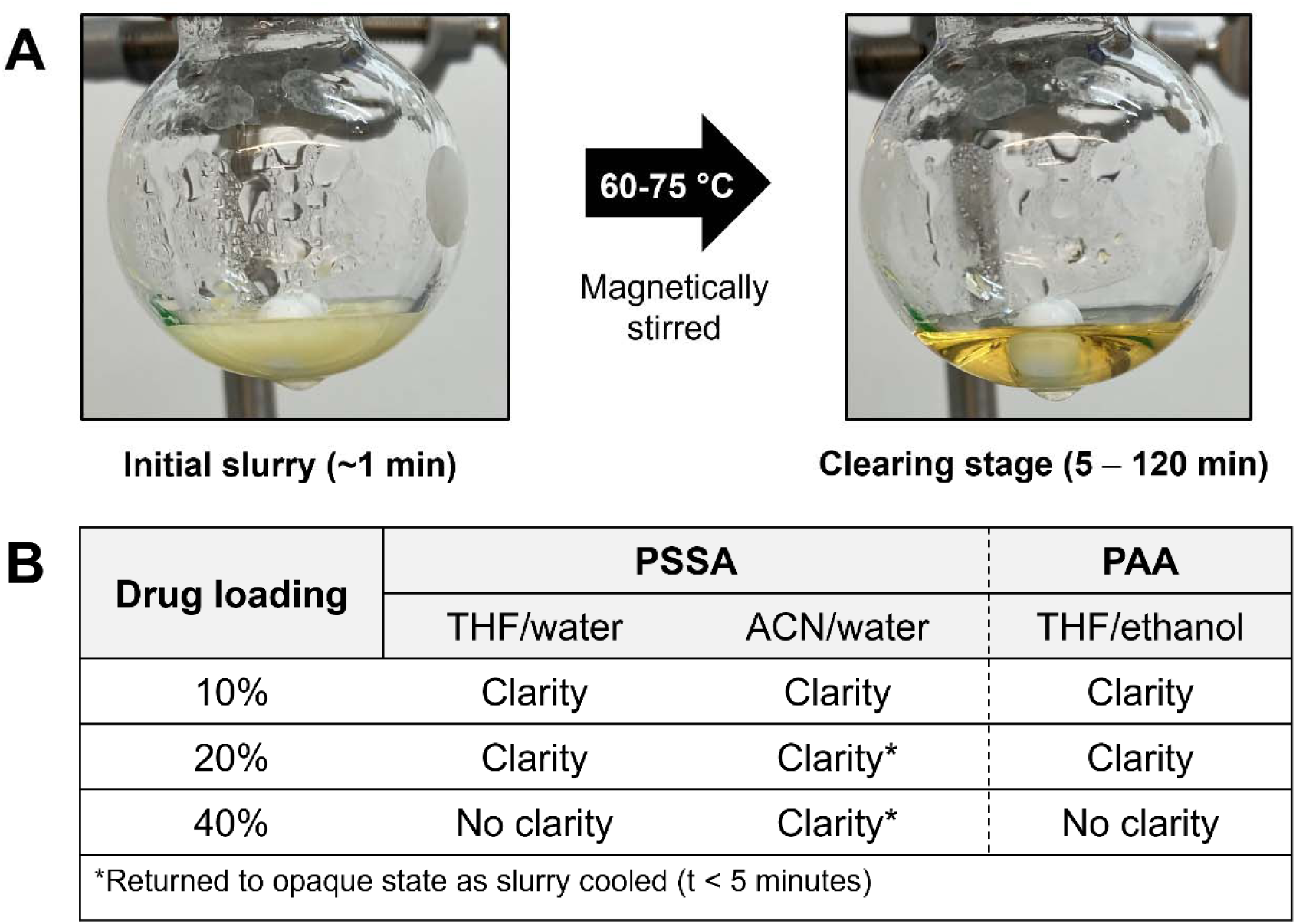
(A) Representative images of slurry formation followed by clearing stage. (B) Table summarizing the slurry clarity for each formulation.

An additional formulation consideration, beyond initial solubilization and achieving clarity, is the energy required to bring the system to dryness. Systems containing a mixed water-solvent environment will require more energy and longer times to dry compared to THF-ethanol systems. The hygroscopic nature of both PSSA and PAA results in water retention within the salt matrix even after initial drying. Dimethyl sulfoxide (DMSO) was also considered as a potential solvent in this work due to the high characteristic solubility of both VBN and PSSA in DMSO, but like water, the elevated boiling point of DMSO (189 °C) would make drying a comparably time-consuming process. Another factor to consider is the dielectric constant, which affects the apparent strength of acids and bases [31]. The accessibility of the reaction sites on the polymers are similarly influenced by solvent type [32].

The THF-water and ACN-water systems provide the necessary conditions for successful complexation. As shown in Figure S.4, after drying the 40% VBN-PSSA (ACN-water) slurry in a vacuum oven, a residual peak corresponding to ACN is observed at 2.05 ppm, indicating incomplete removal of the organic solvent. Upon further drying in a vacuum desiccator, the intensity of this peak decreased significantly. The broad signal centered around ∼5 ppm, which persisted through additional drying, is related to water molecules involved in hydrogen bonding with acidic protons. The continued presence of this peak suggests that residual water is associated with the polymer matrix of the VBN-PSSA salt.

Other drug–polymer slurries in the literature have been described in terms of component mass ratio (drug:polymer, w/w), solvent composition (e.g., ethanol:DCM, v/v), and solids loading (solids:solvent, w/w). We here also report the composition in terms of the charge ratio, i.e. molar ratio of ionizable groups on each species available for complexation during slurry formation. Consistent with prior work, drug polymer slurry formation in this study was conducted using excess acidic groups, up to a 1:56 VBN:PAA charge ratio in the 10:90 w/w formulation [33]. For PSSA formulations, the charge ratio was 1:3 at 40 wt% VBN and 1:20 at 10 wt% VBN. Formulations with a higher excess of acid tended to reach clarity better (**Figure 2B**).

### 3.2 Determining the success of salt formation between the drug and polyanion

VBN is a weakly basic compound with an estimated pK of 2.3. Salt formation between an acid–base pair is favored when the ΔpK exceeds ∼2–3 units, consistent with thermodynamically favorable proton transfer. PSSA (pK ≈ −1.5) exceeds this threshold and is therefore sufficiently acidic to protonate VBN, enabling amorphous drug polymer slurry (ADPS) formation. PAA (pK ≈ 4.5), however, is less acidic than VBN and is not expected to promote proton transfer even when the solvent criteria have been met. To confirm this, the formation of drug–polyanion salts was evaluated using X-ray photoelectron spectroscopy as a surface-sensitive technique to probe elemental composition and chemical state within the near-surface region of materials [34, 35]. In pharmaceutical systems, XPS is used to directly assess nitrogen protonation in amorphous drug–polymer complexes. Protonation of amine groups is typically observed as a positive shift of approximately +2 eV in the N 1s binding energy (BE) [36]. The N 1s spectra for the VBN–polyanion formulations are shown in **Figure 3**.

**Figure 3:**
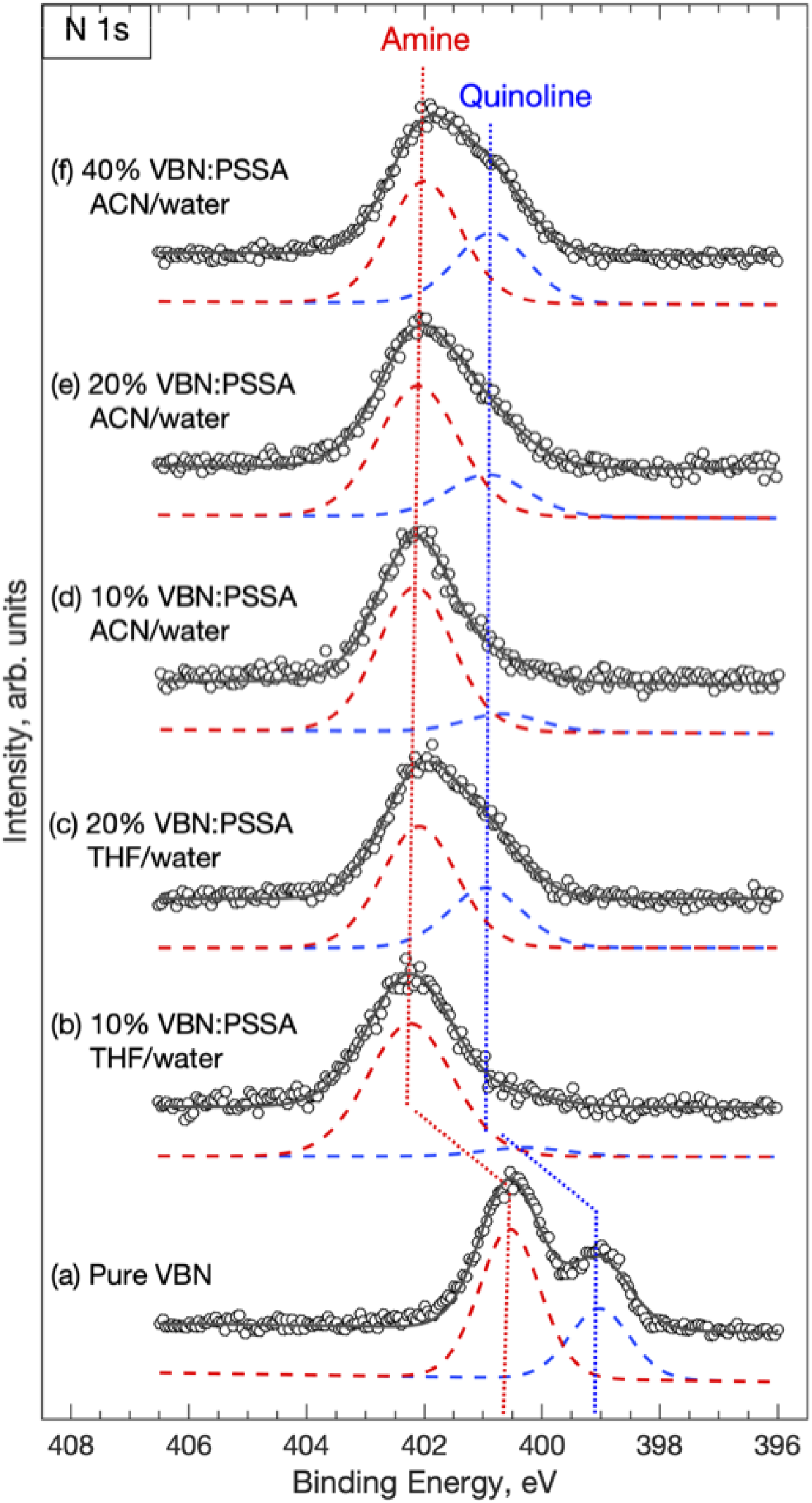
N 1s XPS spectra of pure vodobatinib (A), VBN–PSSA salts formulated with THF/water at 10% (B), and 20% (C), and VBN–PSSA salts formed in ACN/water at 10% (D), 20% (E), and 40% (F) drug loading. Raw data (black circles) were fitted as the sum of two nitrogen-containing component peaks, shown slightly offset for visual clarity.

The N 1s XPS spectra in Figure 3 resolve distinct components corresponding to different nitrogen chemical states within each formulation. Figure 3a exhibits two N 1s peaks at 399.0 and 400.6 eV, which correspond to the quinoline and secondary amine functional groups in crystalline VBN, respectively. All VBN–PSSA ACN/water salts demonstrated peaks of higher BE for both nitrogenous components compared to the pure drug. The relative peak shift range for the secondary amine group and the quinoline nitrogen were 1.5-1.6 eV and 1.6-1.9 eV, respectively (Figure 3d, 3e, 3f). VBN–PSSA THF/water salts show a similar trend, with a quinoline peak shift range of 1.3-1.9 eV and a secondary amine shift of 1.5-1.7 eV (Figure 3b, 3c). Analysis of VBN–PSSA salts across drug loadings and solvent qualities provides insight into the sequence of protonation events. At 10% drug loading (i.e., higher charge ratios; Figure 3b,d), the quinoline nitrogen —being more basic than the amine — is preferentially protonated. However, this protonation appears attenuated in the spectra because the associated shift to higher binding energy overlaps with the crystalline NH signal. As drug loading increases, the NH:quinoline ratio approaches ∼2:1, and both signals are shifted to higher binding energy, indicating that protonation of the amine and quinoline sites occurs to a comparable extent.

Amorphous drug–PSSA salts can experience an electron density redistribution that leads to a positive BE shift for non-protonated components. Similar results were reported for lapatinib–PSSA and gefitinib–PSSA salts showing a 0.6-0.9 eV binding energy increase that was speculated to be due to peak shifts in the F 1s, Cl 2p, and S 2p peak spectra [12]. An analysis of the Cl 2p peaks indicates a <+1.0 eV shift for all VBN–PSSA salts, demonstrating that there is likely a redistribution effect. However, the BE shift observed in **Figure 3** would account for this effect, leaving the remaining change attributed to protonation. Furthermore, N 1s peaks for the 10% VBN–PAA formulation (**SI section S.3**) were closely aligned with pure VBN, indicating that negligible protonation occurred during the slurry manufacture. These results demonstrate that formulations containing PSSA as the polyanion were able to form successful ADPS with VBN, while the formulations containing PAA did not, in line with the proposed hypothesis regarding the pK_a_ values of the respective components.

### 3.3 Characterization of drug amorphous character in the slurry

Amorphous solids tend to exhibit more favorable dissolution kinetics than the crystalline form of the same drug. In the amorphous state, the free energy requirement to remove a molecule from the disordered lattice is lower than that of removing a drug molecule from a highly ordered crystal lattice [11, 37]. For drug–polymer salts, the resulting materials are expected to exhibit amorphous character due to the presence of strong ionic interactions between the drug and polymer, which are typically stronger than the intermolecular forces governing neutral formulations and can enhance resistance to recrystallization. In addition, incorporation into amorphous, water-soluble polymers such as PSSA and PAA reduces the thermodynamic driving force for crystallization, resulting in a more stable amorphous solid product, with prior studies demonstrating that increased salt formation correlates with improved physical stability [17]. Therefore, X-ray diffraction analysis was performed to evaluate the solid-state character of the VBN–PSSA and VBN–PAA formulations (**Figure 4**).

**Figure 4:**
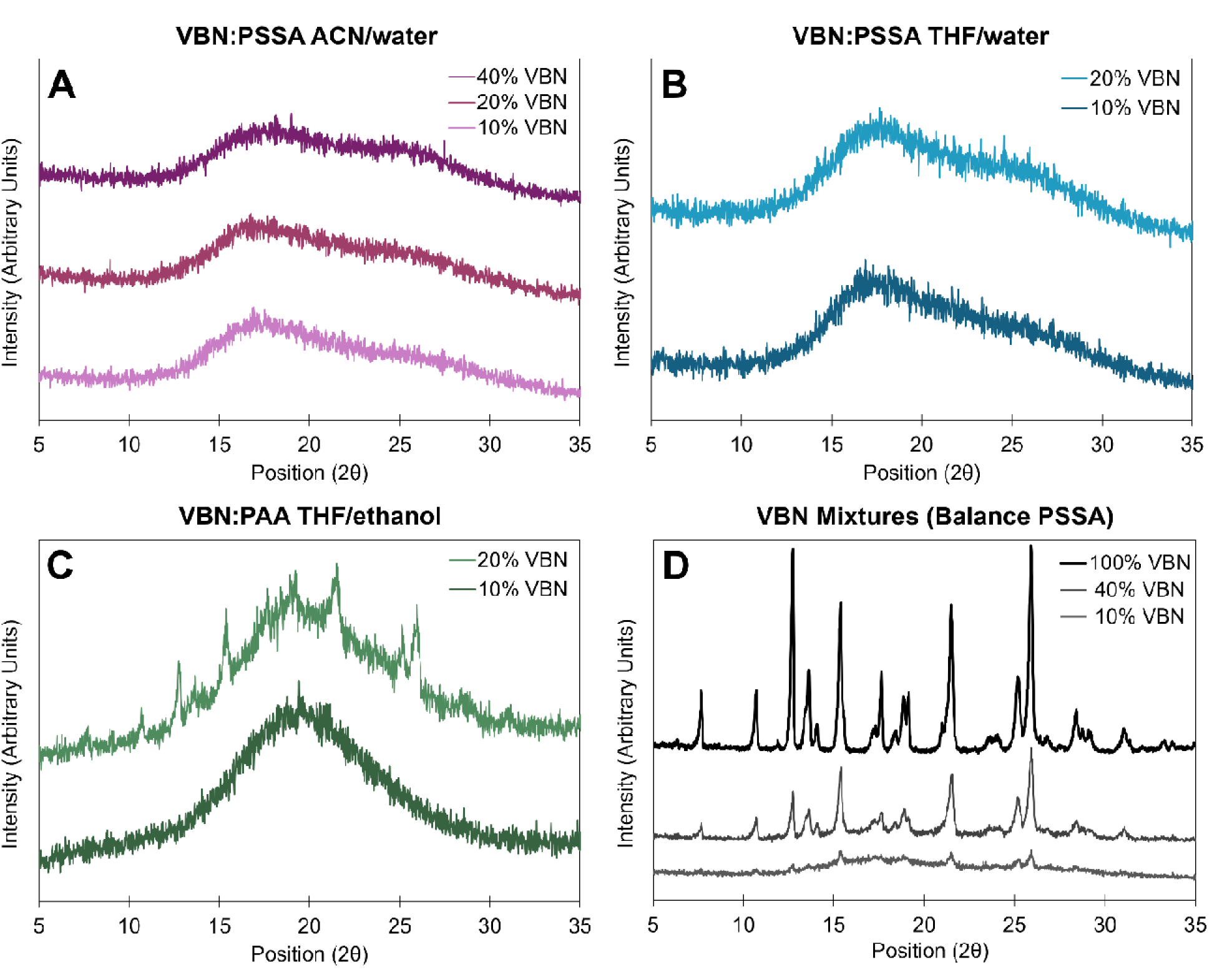
(A-C) Powder X-ray diffraction traces for the drug-polyanion slurries presented herein formulating VBN with PSSA (A&B) and PAA (C). (D) Powder X-ray diffraction analysis for a physical mixture of vodobatinib (VBN) and balance PSSA.

Amorphous drug-polymer salt formulations containing PSSA as the polyanion exhibit amorphous character across all drug weight fractions tested. This is notable because two of these formulations (ACN/water, 20% and 40% VBN) did not maintain prolonged clarity following initial preparation. Since all three ACN/water PSSA-VBN formulations were amorphous, no clear trend can be identified relating amorphous character to drug weight fraction or acid:base charge ratio at this time. The retention of amorphous character despite the loss of visual clarity indicates that clarity is a time-dependent indicator of drug–polyanion salt formation.

Formulations prepared with PAA display measurable crystalline character at 20% drug loading (**Figure 4c**), which is consistent with the XPS analysis that VBN–PAA systems did not exhibit nitrogen protonation and were therefore not stabilized through ionic interactions, but rather through weaker hydrogen bonding. Notably, both the 10% VBN–PSSA and 10% VBN–PAA formulations exhibited amorphous character according to pXRD analysis. This finding, together with the XPS results, indicates that the observed XPS shifts present in VBN-PSSA samples arises from ionic protonation of VBN and is not simply a consequence of the difference between the API in a crystalline versus amorphous state. Overall, the pXRD and XPS results together indicate that visual clarity alone, regardless of timescale, is not a sufficient indicator of successful ADPS formation, and that the magnitude of the pK difference between the drug and polyanion in conjunction with XPS analysis is the most reliable approach to determining ionic complexation.

### 3.4 Dissolution kinetics testing in simulated gastrointestinal fluids

The concentration of VBN in simulated fed-state gastrointestinal fluids was measured over 6 hours from the formulations presented herein in comparison to the crystalline drug to determine if ADPS formulation enhanced dissolution kinetics. For each sample, the area under the curve (AUC) was determined using the trapezoidal rule in Excel. The AUC data obtained from individual replicates are shown in **Figure 5B & 5D**, and the average AUC across replicates of a given formulation are given in **Figure 5E**. Dissolution kinetics curves are presented in **Figure 5A and 5C**.

**Figure 5:**
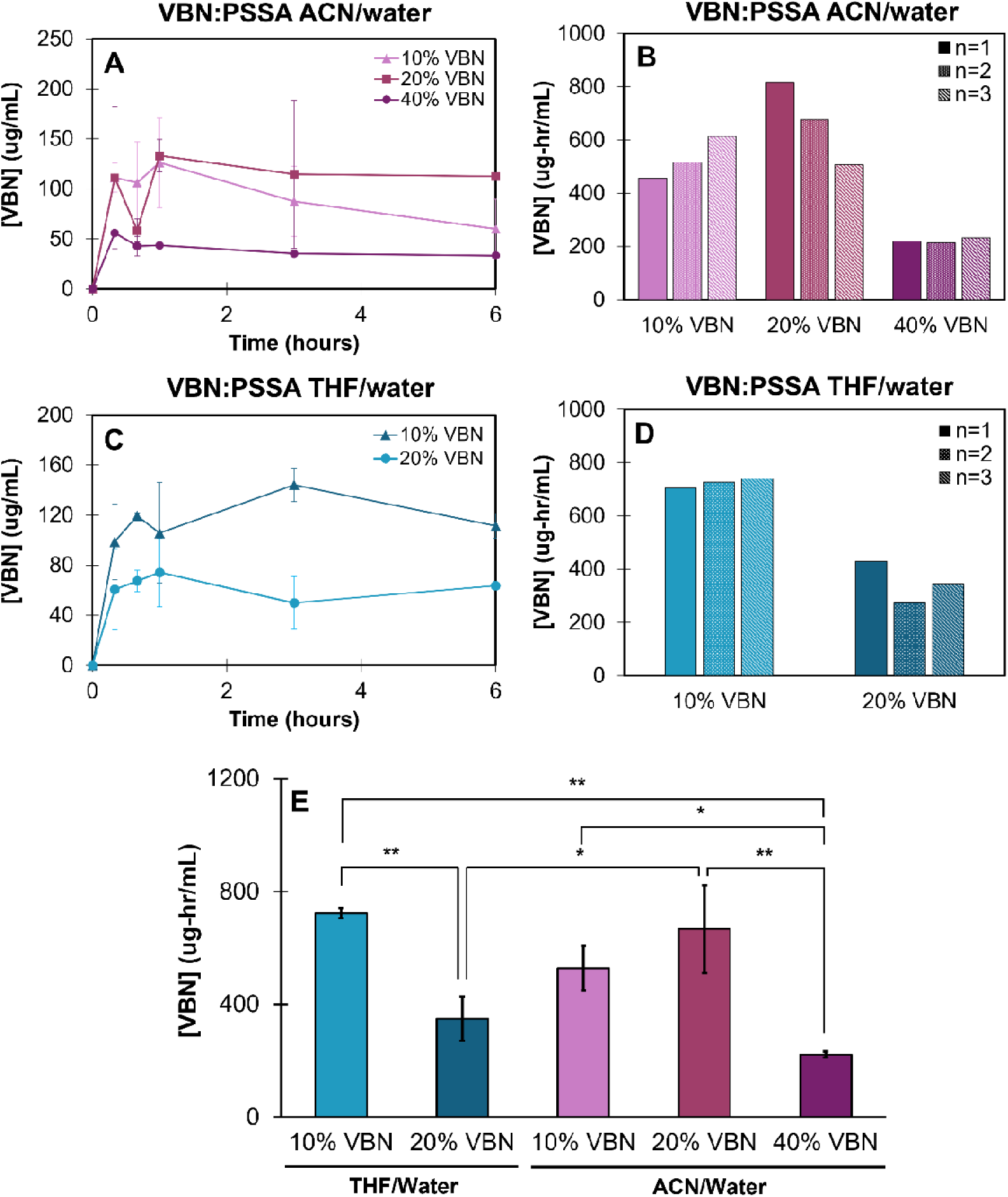
(A) Dissolution kinetics profiles and (B) area under the curve individual replicate analysis for formulations containing VBN and PSSA in ACN/water. (C) Dissolution kinetics profiles and (D) area under the curve individual replicate analysis for formulations containing VBN and PSSA in THF/water. (E) Bar charts representing the average area under the curve over 6 hours for each formulation where the error bars represent the standard deviation of n = 3 release kinetics tests. The display statistics reflect the outcome of a one-way ANOVA with Tukey’s multiple comparisons testing as discussed in the SI (**SI Section S4**). Asterisks correlate with significant p-values following p < 0.01 = **, p < 0.05 = *.

At 10% VBN drug loading, VBN-PSSA slurries formulated using THF exhibited a higher mean AUC than those formulated with ACN (724 ± 15 ug-hr/mL vs 529 ± 65 ug-hr/mL), though the difference was not significant. When the weight fraction of VBN was increased, the trend reversed; at 20% drug loading, VBN-PSSA slurries made with THF (349 ± 64 ug-hr/mL) had a significantly lower AUC than formulations made with ACN (667 ± 127 ug-hr/mL). Within each solvent system, formulations with lower VBN loading exhibited significantly superior AUC compared to more highly loaded formulations, with the exception of 10% vs 20% VBN in PSSA made with ACN-water, which exhibited AUCs similar to one another and each significantly higher than 40% VBN in the same system.

Only formulations containing PSSA as the ionic polymer exhibited quantifiable dissolution kinetics; neither the crystalline drug nor formulations prepared with PAA showed any measurable release under the same conditions. This is consistent with a prior study formulating VBN into polymeric nanocarriers by co-precipitating it with other hydrophobic species (‘co-cores’), in which formulations with higher degrees of crystallinity exhibited slower release kinetics.[30] In that study, the VBN weight fraction in the nanoparticles, which spanned a similar range to the work here, did not strongly affect release kinetics for a suite of formulations with a given co-core, however.

The effect of drug loading on dissolution kinetics has not been directly considered in previous studies of ADPS formation, and the mechanisms governing drug release from these systems remain underexplored [15, 17, 19]. Applying principles developed for standard ASD systems may provide a starting point when considering the relationship between drug loading and dissolution in ADPSs. In ASDs, above a threshold drug loading termed the Limit of Congruency (LoC), a rapid decline in drug release rate occurs, which is attributed to phase separation between drug and polymer upon hydration [38, 39]. Below the LoC, the drug exists as discrete domains dispersed within the polymer matrix that readily dissolve upon hydration, resulting in release rates largely governed by polymer dissolution. At drug loadings above the LoC, these drug domains overlap and eventually form channels that introduce physical and thermodynamic barriers to release. The LoC itself has been shown to be governed by the morphology of the phase separation that occurs upon hydration, which is a function of interactions between the two species [38], [40]. We hypothesize that the strong attractive ion pairing interaction between the drug and polyelectrolyte in ADPSs likely increases the LoC compared to ASDs, by overcoming or at least mitigating drug-drug attractive interactions. The low tendency of drugs in ADPSs to recrystallize, due to low molecular mobility and homogeneous distribution throughout the drug matrix that occur as a result of the electrostatic interaction, possibly supports this idea [19]. Future studies of drug domain size and phase separation morphology in ADPSs as a function of drug loading will be needed to understand how lessons learned for standard ASDs can be applied or adapted for the ADPS system.

## 4. Conclusions

Slurry conversion can be used to generate amorphous drug–polymer salts of vodobatinib, a weakly basic compound with limited enteric solubility. Poly(styrene sulfonic acid) facilitated protonation of the quinoline nitrogen and formation of amorphous salts across multiple drug loadings, confirmed using X-ray photoelectron spectroscopy and powder X-ray diffraction. PSSA has a pKa 3.8 pH units lower than VBN and exhibited salt formation with VBN, while PAA, which has a pKa 2.3 units above VBN, did not. Despite drying successful formulations in a vacuum oven for more than 24 hours, residual solvent was detected by ¹H NMR analysis, indicating a possible need for a secondary drying step.

These findings support slurry conversion as a practical approach to polymeric amorphization strategy for weakly basic compounds and provide a framework for further refinement of this approach to improve dissolution performance. We also recommend reporting drug:polymer charge ratio when discussing the formulation of amorphous drug-polymer slurries. Future work should investigate how the characteristics and release behavior of the amorphous drug–polymer slurries evolve during long-term storage and also explore secondary drying processes for the final product, since residual solvent or moisture may destabilize the amorphous solid and be unacceptable to regulators.

## Acknowledgements

This research was supported by a grant from Sun Pharma Advanced Research Corporation. We thank Dhananjay Umrani, Yashoraj Zala, Minaketan Routray, Shilpa Sule, Saleem Shaikh, Sangeeta Mirgal, Pawan Mishra, Alpana De, and Rajesh Ranganathan for their continued input and guidance throughout the completion of this project.

