## Supporting information for "Improving oral dissolution kinetics of weakly basic vodobatinib via slurry conversion to an amorphous drug-polymer salt"

**S1. Supplemental for materials/methods**

Slurry conversion for each formulation was performed using the experimental setup in Figure S.1. The slurry was mixed inside a clamped round-bottom flask with a magnetic stir bar (mixed at 250 rpm). The apparatus was situated on a 2-in-1 stir magnetic stirrer and heating plate. The flask was semi-submerged in stirred silicone oil that was kept at the desired temperature using a 100 °C thermometer. Due to the highly adhesive properties of both poly(acrylic acid) (PAA) and poly(styrene sulfonic acid) (PSSA), the slurries would be poured into small silicone wells lined with aluminum foil after the clearing stage was reached. After drying, the now solid salt product was transferred into a large ceramic mortar and pestle for grinding with liquid nitrogen. A secondary grinding step was often employed in an agate mortar and pestle for generating finer particles. The resulting powders are shown in Figure S.2.

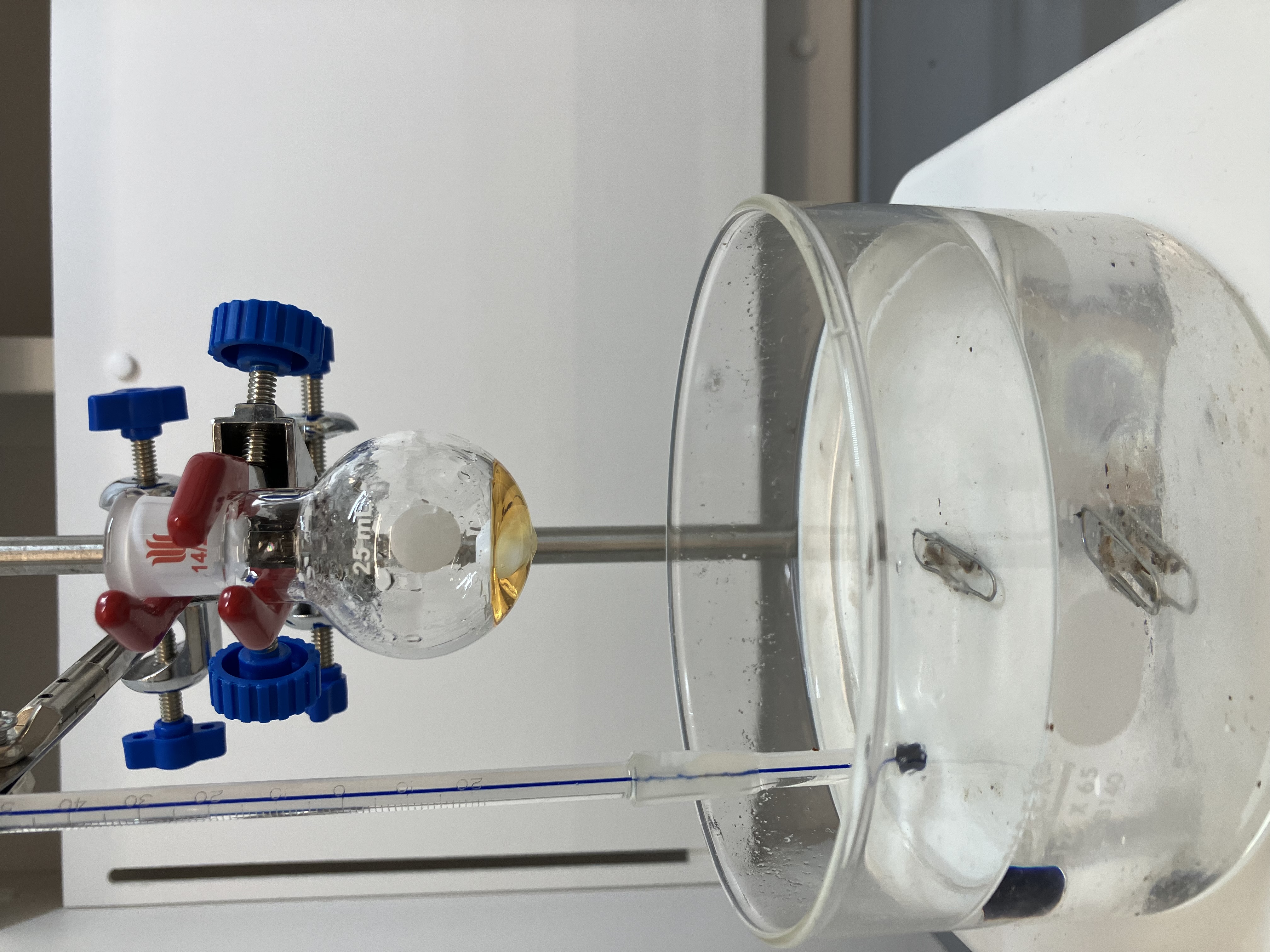

**Figure S.1:** Experimental setup for slurry conversion.

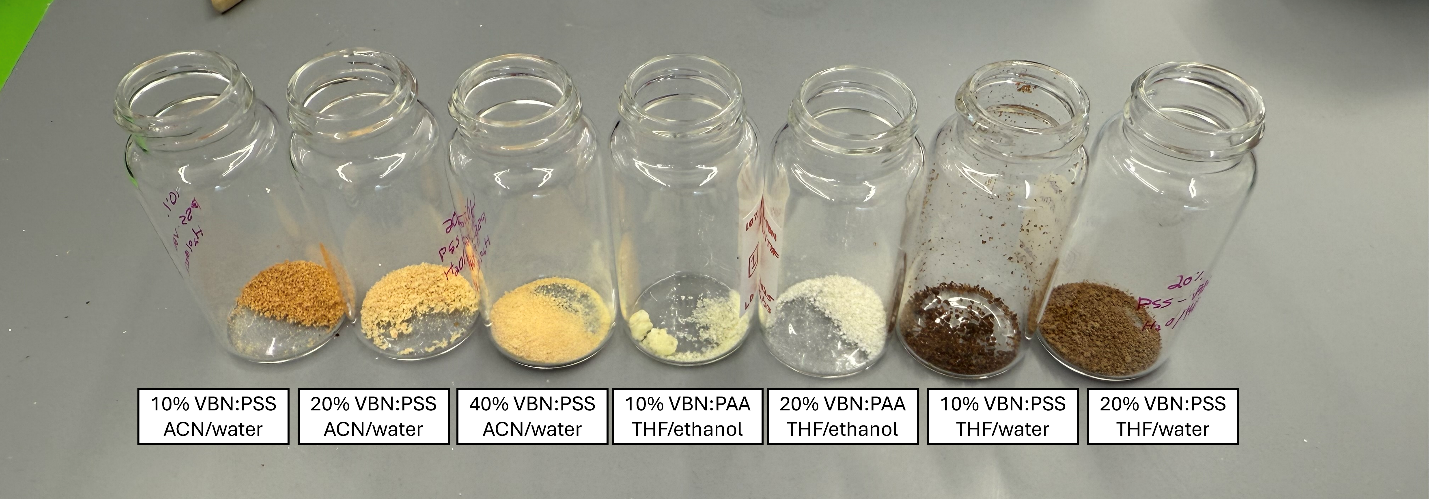

**Figure S.2:** Final powdered form for all drug–polymer formulations.

**S2. Supplemental for 3.1, Slurry clearing**

^1^H NMR data was collected for analyzing residual solvent and indication of chemical reactions between VBN and the corresponding polyanion.

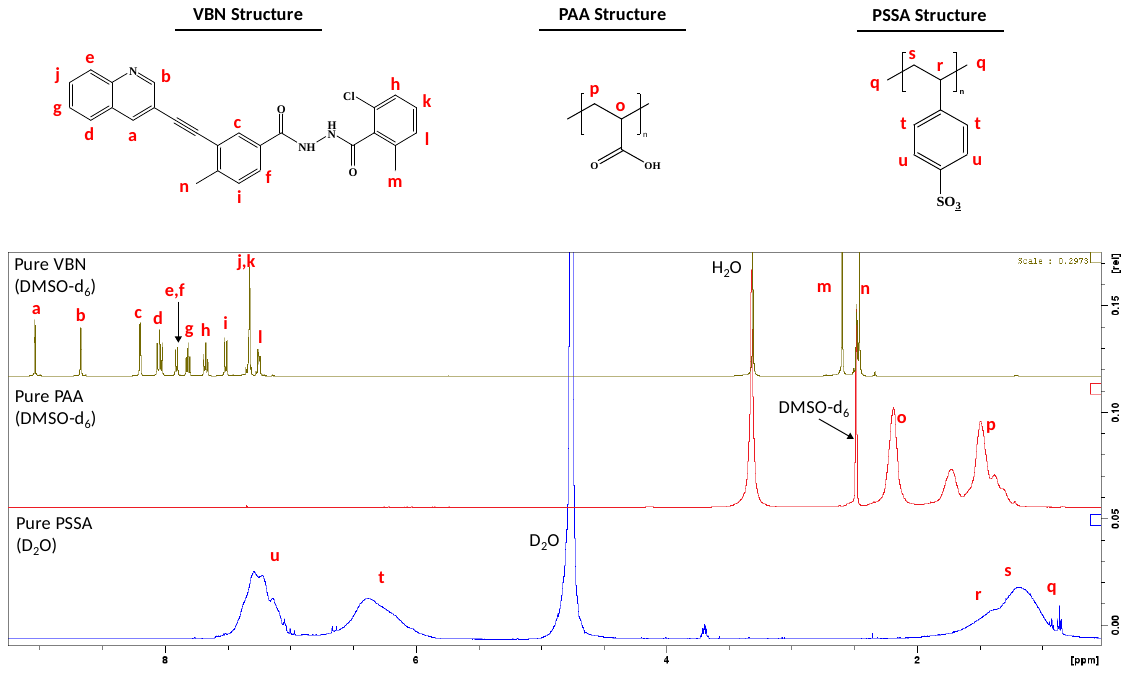

**Figure S.3:** ^1^H NMR Spectra of pure VBN, PAA, and PSSA. DMSO-d_6_ was used as the solvent for VBN and PAA, while D_2_O was used for PSSA.

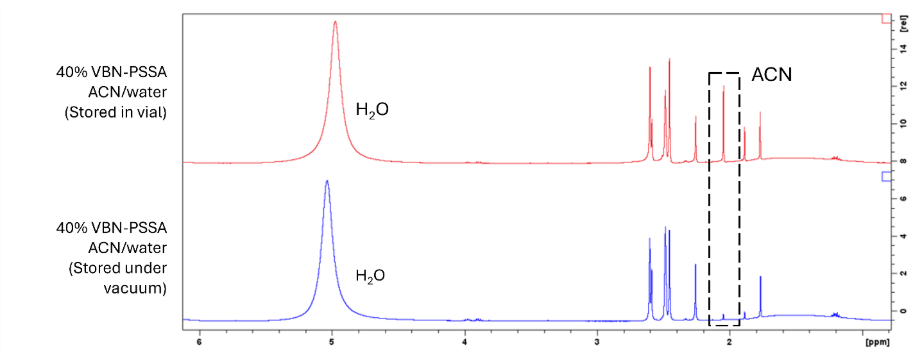

**Figure S.4:** ^1^H NMR Spectra (DMSO-d_6_) of 40% VBN-PSSA in ACN/water dried in vacuum oven and after 24h further drying in a vacuum desiccator.

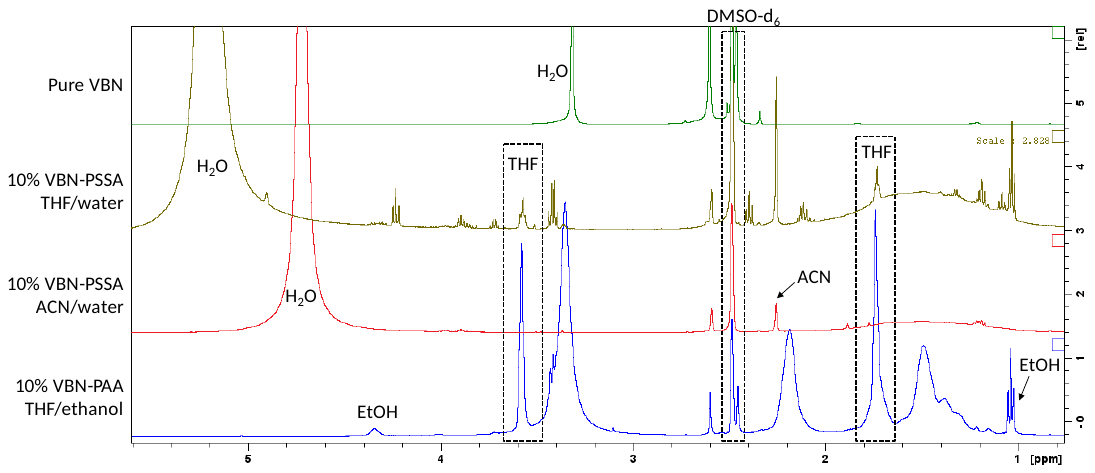

**Figure S.5:** ^1^H NMR Spectra (DMSO-d_6_) comparison of residual solvent in pure VBN and all salt formulations at 10% drug loading. Each salt sample was dried using a vacuum oven for 4 days. Pure VBN was dried in a vacuum desiccator.

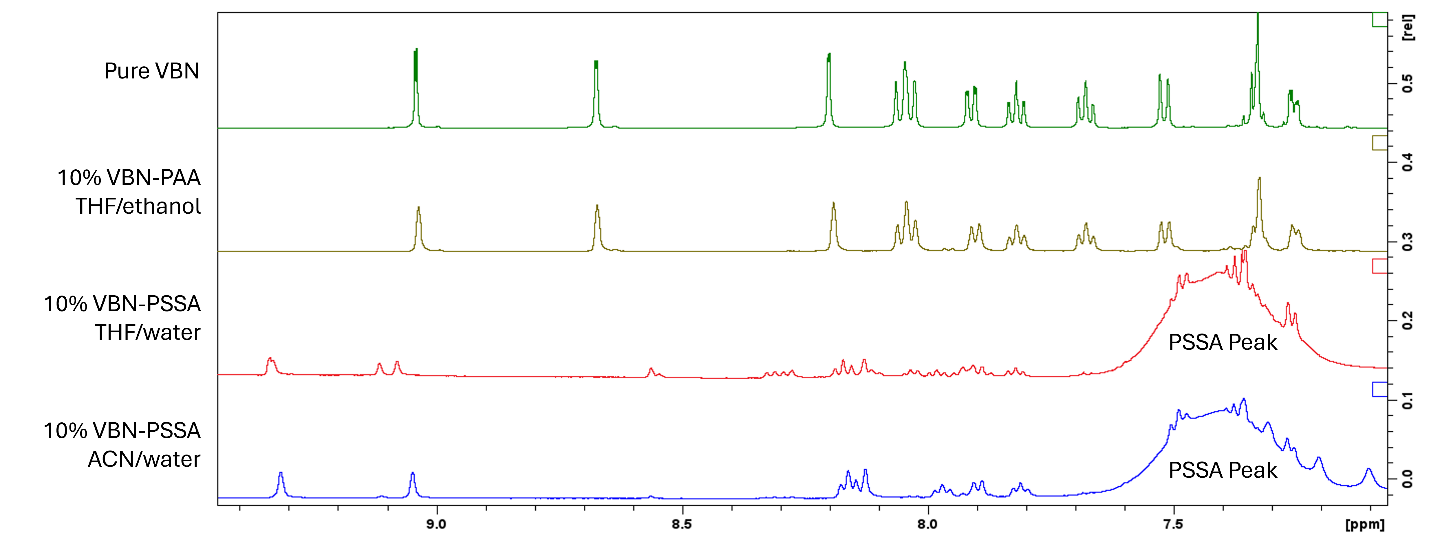

**Figure S.6:** ^1^H NMR Spectra (DMSO-d_6_) comparison of carbon components in pure VBN and all salt formulations at 10% drug loading. Each salt sample was dried using a vacuum oven for 4 days. Pure VBN was dried in a vacuum desiccator.

**S3. Supplemental for 3.2, XPS Analysis**

As describe in the main text, the N 1s peaks determined via XPS analysis for the 10% VBN–PAA formulation were closely aligned with pure VBN, indicating that no protonation occurred. This result is presented in Figure S.7 and is expected due to the difference in pKa’s between the two molecules. The 20% VBN–PAA formulation was not analyzed due to its apparent crystallinity as found through PXRD.

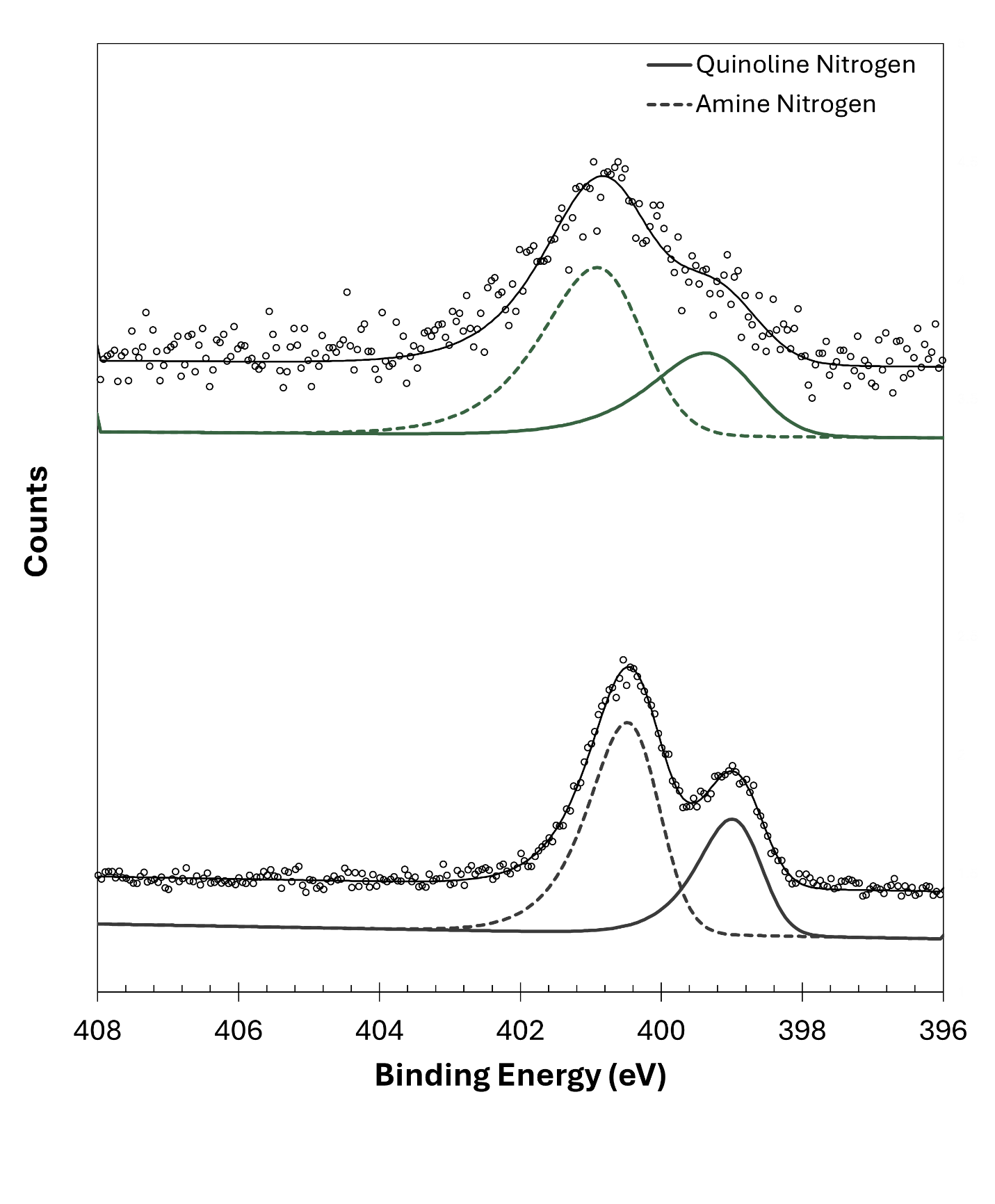

**B**

**A**

**Figure S.7**: N 1s XPS spectra of (A) 10% drug loaded VBN–PAA salt formulated in THF/ethanol and (B) pure vodobatinib. Raw data (black circles) were fitted as the sum of two nitrogen-containing component peaks, shown slightly offset for visual clarity.

C 1s peak fitting for charge correction is shown in Figure S.8. Relevant carbonyl groups were fitted to determine the peak binding energy of the aliphatic carbon (C–C), which was then manually aligned to 284.8 eV

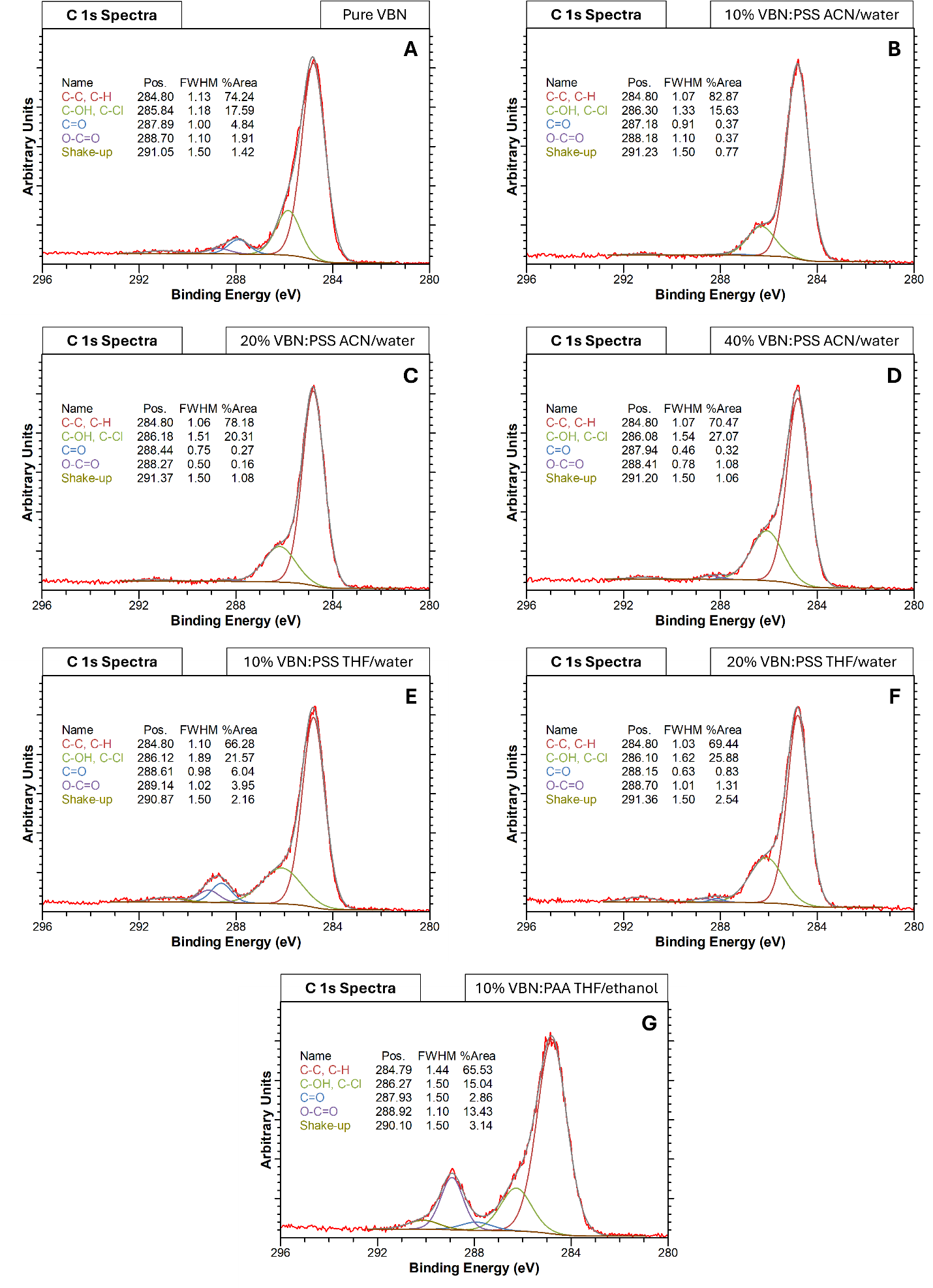

**Figure S.8**: C 1s XPS spectra of (A) pure VBN, (B-D) VBN–PSSA salts formulated with ACN/water at 10, 20, and 40% drug loading, (E-F) VBN–PSSA salts formulated with THF/water at 10 and 20% drug loading, and (G) VBN–PAA formulated with THF/ethanol at 10% drug loaded.

**S4. Supplemental for 3.4, Dissolution kinetics testing**

S4.1 Statistical analysis of area under the curve data

Statistical analysis of the AUC data was performed using a one-way ANOVA with Tukey multiple comparisons testing. This calculation was supported by [www.BioRender.coms/](http://www.BioRender.coms/) graphing feature. The tabulated results of the analysis are provided below.

**Table S.1:** Test summary for a one-way ANOVA with Tukey’s multiple comparisons.

| **Comparison** | **Adj. P-value** | **ΔMeans** | **95% CI of Δmeans** |
| --- | --- | --- | --- |
| THF/MQ 10% vs 20% | 0.007993** | 384.24 | 106.4843 to 661.9901 |
| THF/MQ 10% vs ACN/MQ 10% | 0.1784ns | 204.99 | -72.7597 to 482.7460 |
| THF/MQ 10% vs ACN/MQ 20% | 0.9222ns | 66.62 | -211.1351 to 344.3706 |
| THF/MQ 10% vs ACN/MQ 40% | 0.001127** | 511.73 | 233.9759 to 789.4816 |
| THF/MQ 20% vs ACN/MQ 10% | 0.1927ns | -179.24 | -427.6738 to 69.1857 |
| THF/MQ 20% vs ACN/MQ 20% | 0.01301* | -317.62 | -566.0492 to -69.1897 |
| THF/MQ 20% vs ACN/MQ 40% | 0.4657ns | 127.49 | -120.9382 to 375.9213 |
| ACN/MQ 10% vs ACN/MQ 20% | 0.3938ns | -138.38 | -386.8051 to 110.0543 |
| ACN/MQ 10% vs ACN/MQ 40% | 0.01600* | 306.74 | 58.3059 to 555.1653 |
| ACN/MQ 20% vs ACN/MQ 40% | 0.0013710** | 445.11 | 196.6813 to 693.5407 |
